# Leiomodin 2 functions as a processive pointed-end elongator of actin filaments

**DOI:** 10.64898/2026.05.04.720848

**Authors:** Sudipta Biswas, Tania M. Larrinaga, Sandeep Choubey, Carol C. Gregorio, Shashank Shekhar

**Author notes:** **Correspondence**: Shashank Shekhar.

## Abstract

The actin cytoskeleton drives essential processes like cell migration and muscle contraction. While barbed-end polymerization is well-established, pointed-end elongation was long considered impossible *in vivo*. Here, we demonstrate that Leiomodin 2 (Lmod2), which localizes to thin-filament pointed ends (PEs) in striated muscle cells, functions as the first identified eukaryotic processive actin polymerase. Single-molecule and single-filament imaging reveal that Lmod2 stably associates with PEs *in vitro*, enabling elongation even in the presence of high profilin concentrations found in the cytoplasm that otherwise would cause depolymerization of free PEs. We find that both processivity and elongation rate of Lmod are dependent on its WH2 domain. Remarkably, human dilated cardiomyopathy-associated mutations in Lmod2 greatly reduce Lmod2’s PE elongation activity, providing a potential mechanism for disease progression, underscoring the essential role of its actin polymerase activity in formation and maintenance of muscle sarcomeres.

## INTRODUCTION

Vertebrates rely on their actin cytoskeleton to generate forces required for essential processes like cell migration, wound healing, and muscle contraction^1,2^. Cellular actin networks are thought to assemble by the addition of ATP-actin monomers at filament barbed ends (BEs), while depolymerization is thought to occur primarily at filament pointed ends (PEs) by dissociation of ADP-actin monomers^2^. This asymmetry between filament ends is reinforced by numerous actin-binding proteins in cells. Profilin-bound monomeric actin (G-actin), the predominant polymerizable actin species in cells, binds exclusively to BEs but not PEs of pure actin filaments^3,4^, restricting elongation to this end. This asymmetry is further amplified by formins which promote BE end polymerization from profilin-actin^5,6^. At the same time, cofilin and cyclase-associated protein drive rapid PE depolymerization^7-10^. Although mechanisms promoting BE disassembly have been identified^11,12^, no eukaryotic protein has been shown to processively polymerize actin filaments from their PEs^13^.

This gap in our understanding is particularly evident in striated muscle cells, where actin “thin” filaments and myosin “thick” filaments organize into sarcomeres, the basic contractile units of striated muscle^14^. CapZ tightly caps the BEs of sarcomeric actin filaments (*K*_*D*_ = 0.1 nM) at the Z disc to prevent monomer addition or removal^15^. Yet, these filaments can elongate, as demonstrated by studies in isolated cardiomyocytes where monomer addition was surprisingly observed at PEs^16^. However, the molecular mechanism that enables PE elongation remains unknown.

Lmod2 is a member of the tropomodulin (Tmod) protein family^17^, which localizes near the PEs of thin filaments in skeletal and cardiac muscle cells. Lmod2 and Tmod1 are together essential for actin filament length regulation in striated muscle cells^16,18-20^. While Lmods have been shown to promote actin nucleation *in vitro*^21^, this activity alone cannot fully account for their *in vivo* roles. Loss of Lmod2 in isolated murine cardiomyocytes, knockout models, and iPSC-derived human cardiomyocytes results in thin filament shortening^22-26^, whereas overexpression leads to aberrant elongation^27^, supporting the hypothesis that Lmod2 acts as an actin filament elongation factor. However, direct molecular evidence for its polymerase activity has thus far been lacking. Here, we show that Lmod2 is a processive actin polymerase that directly elongates actin filaments from their PEs.

Apart from Lmod2, vertebrates express two additional Lmod isoforms (Lmod1 and Lmod3), each encoded by distinct genes essential for muscle function. Mutations in Lmod isoforms cause severe muscle diseases. *LMOD1*, the predominant smooth-muscle isoform, is mutated in megacystis microcolon intestinal hypoperistalsis syndrome^28^. *LMOD2*, the predominant cardiac isoform, is mutated in dilated cardiomyopathy (DCM), with homozygous loss-of-function variants such as W398* causing severe neonatal DCM requiring heart transplantation^23,29-32^. Mutations in *LMOD3*, the main skeletal-muscle isoform, cause nemaline myopathy (NM)^33-35^.

To elucidate Lmod2’s role in actin dynamics, we used microfluidics-assisted total internal reflection fluorescence (mf-TIRF) microscopy to directly observe its effects on actin filament assembly. We found that, beyond nucleating filaments, Lmod2 can also polymerize actin filaments at their PEs in the presence of profilin-bound G-actin, a condition in which free PEs depolymerize. Using multispectral single-molecule imaging, we directly visualized individual Lmod2 molecules bound to filament ends, remaining processively associated for several minutes. Lastly, we show that Lmod2’s regulation of filament length depends on its WH2 domain, which controls both polymerization rate and processivity.

## RESULTS

### Lmod2 remains processively bound to elongating PEs

To investigate Lmod2’s role in actin assembly, we immobilized biotinylated SNAP-tagged mouse full-length Lmod2 on streptavidin-coated, PEG-silane–passivated coverslip within an mf-TIRF chamber^36^ (Fig. 1A,B). Upon introducing Alexa-568–labeled (red) G-actin, newly nucleated filaments appeared and remained anchored to the coverslip through Lmod2 (Fig. 1C). In control experiments, without Lmod2, no coverslip-attached filaments were observed (movie S1, Fig. S1). These results confirm Lmod2’s established nucleation activity^21^ and further suggest that it can remain bound to newly nucleated filaments.

**Fig. 1:**
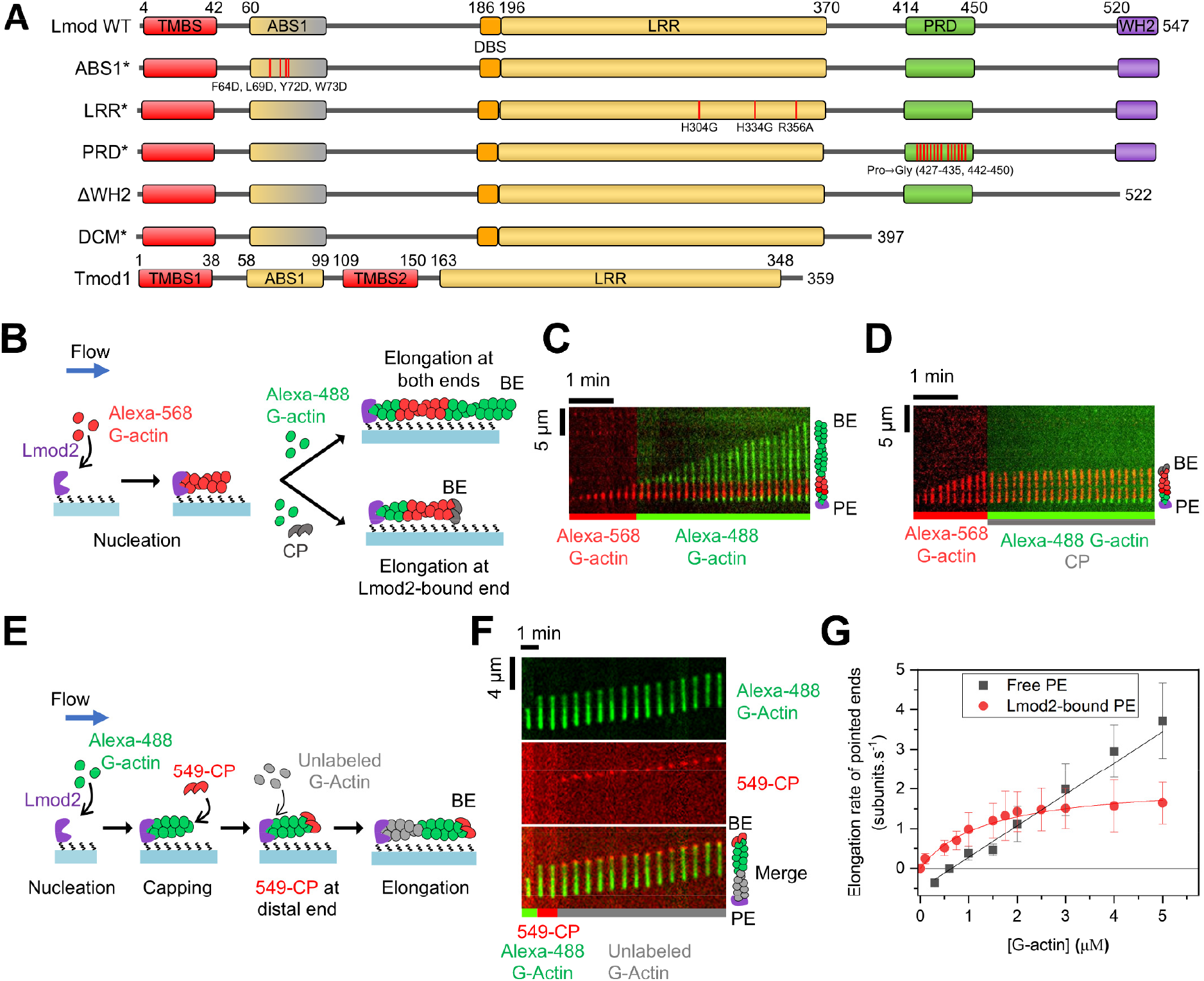
Lmod2 drives processive PE polymerization. **(A)** Domain diagrams of human Lmod2 wildtype, mutants and deletions used in this study. Tmod1 domain diagram is shown for comparison. TPMBS, Tpm-binding site; ABS1, actin-binding site-1; DLBS, D-loop–binding site; LRR, Leu-rich repeat domain; PRD, proline-rich domain; WH2, WASP homology-2 domain. Mutated amino acids are indicated. ABS1* carries four mutations in the region corresponding to the first actin-binding site of Tmod1. While the N-terminal region of ABS1 shows partial conservation in Lmod2, the C-terminal site does not, as indicated by the gradient coloring. In PRD*, all proline residues in the ranges 427–435 and 442–450 were replaced with glycine. **(B)** Experimental strategy: filaments nucleated from coverslip-anchored Lmod2 and 1 µM red G-actin (15% Alexa-568), elongating from 2 µM green G-actin (15% Alexa-488) ± 25 nM capping protein (CP). **(C–D)** Representative kymographs of filaments elongating under these conditions without (Movie S2) or with 25 nM CP (Movies S3-S4). **(E)** Experimental strategy: filaments nucleated from coverslip-anchored Lmod2 and 1 µM green G-actin (15% Alexa-488), transiently exposed to 10 nM 549-CP (red), and then elongating from 2 µM unlabeled G-actin. **(F)** Representative kymograph of a filament (top) capped by 549-CP (middle) elongating under these conditions (bottom, merged; Movie S5). **(G)** Elongation rates (mean ± SD) of free (black) and Lmod2-bound (red) PEs at varying G-actin concentrations. Symbols, experimental data; lines, linear fit (black) to free PEs data yielding *k*_*+1*_ = 0.79 ± 0.07 subunits.µM.s^−1^ and *k*_*−1*_ = 0.51 ± 0.06 subunits.s^−1^, or to saturating binding curve (red) to Lmod2-bound PE data used to estimate the plateau rate of ∼2.2 subunits·s^−1^. Filament numbers: Lmod2-bound, 32–57 per condition (8 at 0.1 µM); free, 30–70 per condition (10 at 10 µM, 12 at 0.1 µM).

To determine whether actin monomers were added at the Lmod2-bound PE, we introduced Alexa-488–labeled (green) G-actin into the flow chamber following nucleation by Lmod2. Three main observations were made (Fig. 1C, movie S2): (1) the Lmod2-nucleated, red-labeled segment gradually translocated in the direction of the flow; (2) green-labeled actin was added at the exposed BE at a rate of ∼18 ± 2.1 subunits·s^−1^, and (3) most notably, green-labeled actin was also incorporated between the anchored PE and the red-labeled segment at a rate of ∼1.4 ± 0.2 subunits·s^−1^ (2 µM G-actin). To confirm filament polarity, we repeated the experiment in the presence of CP, which specifically binds the BE and inhibits subunit exchange at the BE ^37^. Under these conditions, distal-end elongation ceased, whereas elongation at the Lmod2-anchored PE persisted at the same rate (Fig. 1D, movies S3-S4).

In a different experimental configuration, SNAP-tagged CP (SNAP-CP), labeled with a benzylguanine-conjugated, green-excitable dye (549-CP), served as a BE marker (Fig. 1E). When green filaments nucleated from anchored Lmod2 were exposed to unlabeled G-actin and 549-CP in the flow, the Lmod2-bound PE continued to elongate, while 549-CP was bound to the distal BE of the filament (Fig. 1F). Importantly, 549-CP and the green filament segment moved together at the same speed away from the tethering site, in the direction of flow (Fig. 1F, movie S5). Together, these observations confirm that Lmod2 remains processively associated with the elongating PE after nucleation.

We next asked whether elongation at Lmod-bound PEs depends on G-actin concentration, as it does for free PEs. To measure elongation rates of Lmod-bound PEs, we used the strategy shown in Fig. 1E. Elongation rates of free PEs were determined by exposing actin filaments anchored to coverslip-bound biotin-CP to labeled G-actin (no filaments were captured on the coverslip in the absence of coverslip-anchored biotin-CP; movie S6). Free pointed ends elongated at rates proportional to the actin monomer concentration. The slope of the plot gives the association rate constant *k*_+1_ = 0.79 ± 0.07 subunits.µM.s^−1^ and the Y-intercept gives the dissociation rate constant *k*_−1_ = 0.51 ± 0.06 s^−1^, both similar to previous measurements^38,39^. In contrast, the elongation rate of Lmod-bound PEs increased with actin monomer concentration only at low actin concentrations and approached a plateau of ∼2.2 ± 0.2 subunits·s^−1^ at higher actin concentrations. This behavior is consistent with a reversible second-order association reaction linked to a reversible first-order transition that becomes rate-limiting at high actin concentrations. Note that the individual values of *k*_+2_ and *k*_−2_ (forward and reverse rates of the first order transition) cannot be determined separately from these data alone. This rate-limiting transition may reflect a process associated with Lmod2 tracking the growing PE, such as a conformational transition or stepping reaction, analogous to gating steps previously described for formins^40^.

Interestingly, we observed that at G-actin concentrations below ∼2 µM, Lmod-bound PEs elongated faster than free PEs (Fig. 1G). Importantly, Lmod-bound PEs elongated even at G-actin concentrations below the critical concentration for the free PE (0.6 µM), i.e., under conditions where the free PE depolymerizes. Linear fits restricted to this low-concentration regime yielded a slope, with *k*_+1_ = 0.84 ± 0.08 subunits·µM^−1^·s^−1^, comparable to the G-actin association rate constant measured here for free PE (Fig. 1G) or reported previously ^38,39^. Importantly, the y-intercept of the fit was close to zero (i.e., *k*_*−1*_ ≈ 0), indicating that Lmod binding strongly suppresses subunit dissociation from the pointed end. This is consistent with our experimental observation that Lmod-bound PEs do not exhibit any measurable depolymerization in the absence of free G-actin. Thus, enhanced net elongation of Lmod-bound PEs at low actin concentrations arises primarily from suppression of subunit dissociation rather than an increase in the association rate. Similar behavior was also observed with human Lmod2 (Fig. S2).

### Direct visualization of Lmod2 on elongating PEs

To directly visualize Lmod2 during elongation, we used 549-Lmod2, prepared as described above for 549-CP. Photobleaching analysis revealed that ∼90% of coverslip-adsorbed 549-Lmod2 photobleached in a single step, confirming the monomeric nature of the labeled protein (Fig. S3A,B). A solution containing 549-Lmod2 and green G-actin was first introduced into the mf-TIRF chamber, and filaments were captured by coverslip-anchored biotin-CP (Fig. 2A). Two distinct populations of filaments were observed: those lacking 549-Lmod2 (free PEs) and those having 549-Lmod2 at their PE (18%, 54 of 311). These filaments were then exposed to a flow containing 2 µM labeled G-actin and unlabeled CP. Filament-bound 549-Lmod2 processively tracked the elongating PE (Fig. 2B) (movies S7-S8). Consistent with our observations with unlabeled Lmod2 (Fig. 1G), 549-Lmod2-bound PEs elongated faster than free PEs (Fig. 2C). Importantly, Lmod2 was never observed on filament sides, indicating that it binds exclusively at or near the PE. These findings directly demonstrate that Lmod2 remains bound to and drives processive elongation at the PE.

**Fig. 2:**
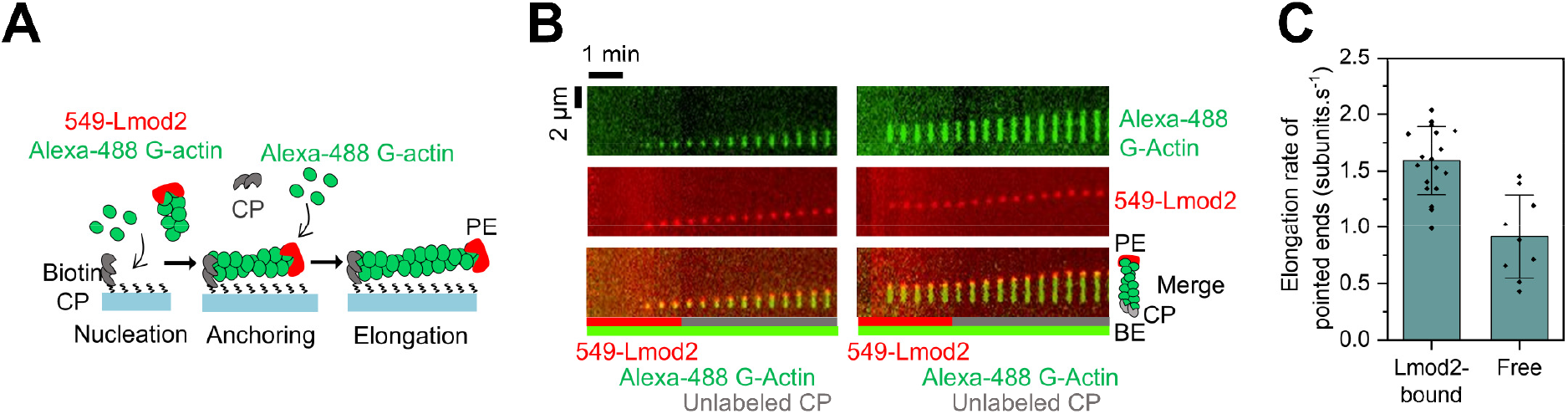
Direct visualization of Lmod2 during PE elongation. **(A)** Experimental strategy: a solution containing 10 nM 549-Lmod2 (red) and 2 µM 15% Alexa-488 G-actin (green) was first introduced into the mf-TIRF chamber, and filaments were captured by coverslip-anchored biotin-CP. These filaments were then exposed to a flow containing 2 µM 15% Alexa-488 labeled G-actin and 25 nM unlabeled CP in TIRF buffer. **(B)** Representative kymographs of two different filaments (top) with 549-Lmod2 bound at their distal PE (middle) elongating under these conditions (bottom, merged; Movies S7-S8). **(C)** Elongation rates (mean ± SD) of PEs for free and 549-Lmod2 bound ends in the presence of 2 µM 15% Alexa-488 G-actin.

### Elongation of Lmod-bound PEs persists in the presence of profilin-actin and Tropomyosin

Profilin binds to the hydrophobic cleft (H-cleft) at the BE of G-actin. Because the H-cleft mediates actin monomer binding at the PE, profilin inhibits this interaction, whereas binding at the BE involves the DNase I-binding loop (D-loop) on the opposite side of the monomer, which is unaffected by profilin. As a result, free PEs depolymerize in the presence of profilin–actin, the primary pool of polymerization-competent monomers in cells, while free BE elongation remains mostly unaffected^3^. We asked whether Lmod2 could utilize profilin-actin as a monomer source for PE elongation, similar to how formins use it at the BE^5,41^. In formins, a PRD mediates the recruitment of profilin-actin, so we further asked whether Lmod2’s PRD contributes to processive PE elongation from profilin-actin.

Lmod2-nucleated filaments continued to elongate from the PE in the presence of equimolar profilin and G-actin, though their elongation rates were reduced compared to filaments grown with G-actin alone (Fig. 3A–C, movie S9). At low actin concentrations (< 2 µM) elongation rates increased approximately linearly with actin concentration, whereas at actin concentrations above 2 µM, elongation rates saturated at ∼1.5 subunits·s^−1^, lower than the plateau observed in the absence of profilin (Fig. 1G). When profilin was added in excess to G-actin, ensuring that >97% of the actin monomers were bound to profilin, a similar qualitative dependence on total actin concentration was observed, but the elongation rate at saturation decreased further (Fig. 3C, S4). This reduced plateau suggests that profilin influences the actin-independent, rate-limiting step in elongation possibly by perturbing Lmod2-dependent transitions (e.g. Lmod2’s stepping) at the pointed end, thereby slowing this step. However, the molecular basis of this effect remains unclear. Importantly, under both conditions free PEs depolymerized, whereas Lmod2-bound PEs continued to elongate (Fig. 3C), demonstrating Lmod2’s ability to promote polymerization under cell-mimicking conditions that typically favor depolymerization. Surprisingly, replacing most proline residues in Lmod2’s PRD region (PRD*, Fig. 1A) had no effect on elongation from profilin-actin, suggesting this domain is dispensable for this activity (Fig. 3D).

**Fig. 3:**
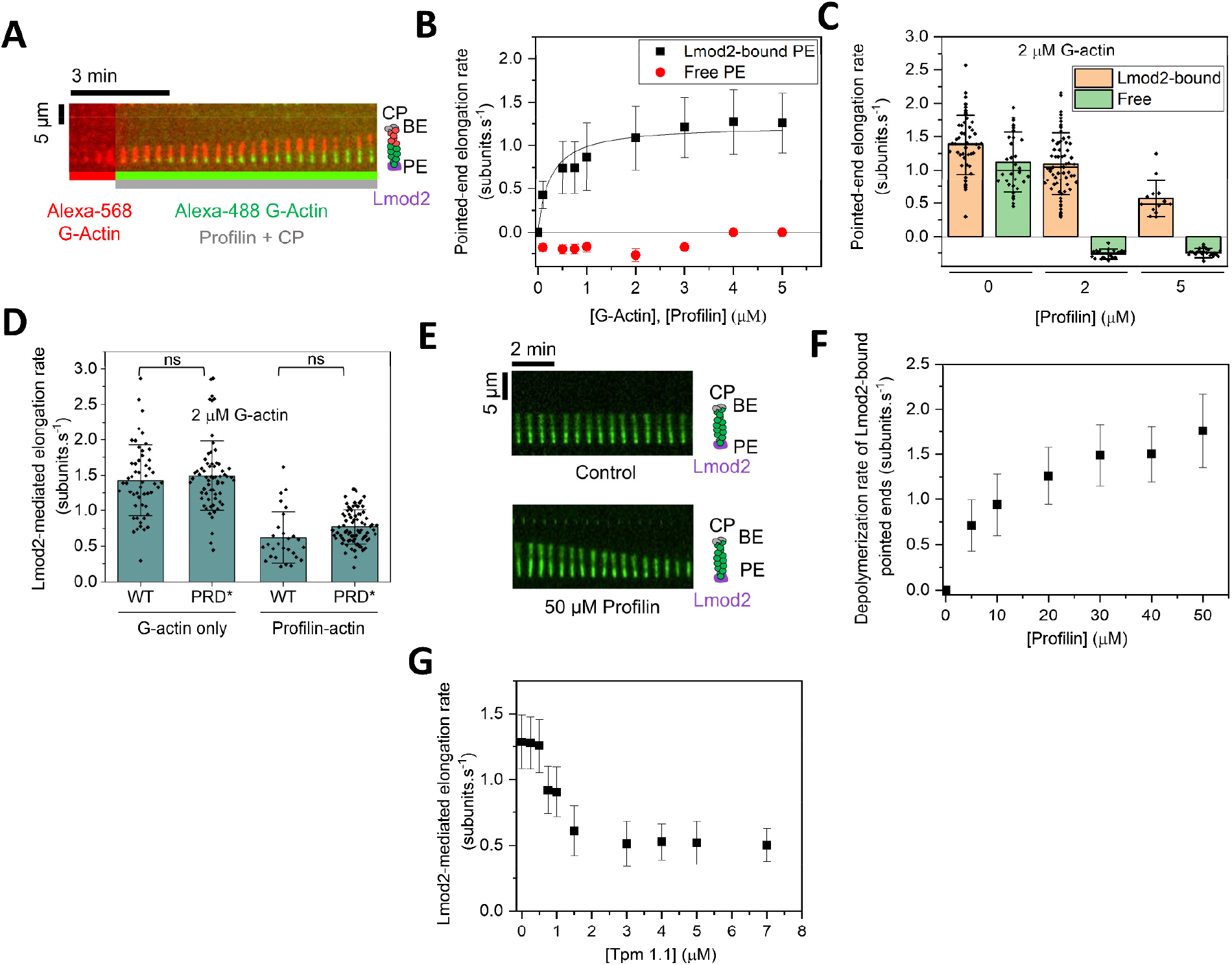
Lmod2-mediated PE elongation persists in presence of profilin and Tpm. **(A)** Representative kymograph of an Alexa-568 (red) filament nucleated by coverslip-anchored Lmod2, and elongating from 2 µM Alexa-488 (green) G-actin and 2 µM profilin (Movie S9). **(B)** Elongation rates (mean ± SD) of Lmod2-bound (black symbols) and free (red symbols) PEs at varying equimolar concentrations of G-actin and profilin. Black line, saturating binding curve used to estimate the plateau rate of ∼1.2 subunits·s^−1^. Number of filaments: Lmod2-bound, 22–71 per condition; free, 20–55 per condition (except 12 filaments at 3 µM). **(C)** Elongation rates (mean ± SD) of Lmod2-bound (orange bars) and free (green bars) PEs with 2 µM G-actin and 0 (left), 2 (middle), or 5 µM profilin (right). Number of filaments (left to right): 44, 40, 71, 25, 62, 65. Also see Fig. S4. **(D)** PE elongation rates (mean ± SD) with Lmod2 WT and PRD mutant, 2 µM G-actin, and without (left) or with 5 µM profilin (right). Number of filaments (left to right): 53, 64, 28, 80. Statistical comparison by two-sample t test against WT (ns, p < 0.05). **(E)** Representative kymographs of coverslip-anchored Lmod2-nucleated green filaments capped at the BE with 25 nM CP, depolymerizing in the absence (top) or presence (bottom) of 50 µM profilin (Movie S10). **(F)**Depolymerization rates (mean ± SD) of Lmod2-bound PEs as a function of profilin concentration in the presence of 25 nM CP. Number of filaments: 38–60 per condition. **(G)**Elongation rates (mean ± SD) of Lmod2-bound PEs as a function of Tpm1.1 concentration with 1 µM G-actin and 25 nM CP. Number of filaments: 47–58 per condition.

To explore the reason behind reduced elongation with profilin-actin, we examined profilin’s effect (without G-actin) on the length of filaments anchored via Lmod2 and capped at their BE by CP. Surprisingly, filaments depolymerized under these conditions, with the depolymerization rate scaling with profilin concentration (Fig. 3E,F). Fiduciary marker tracking confirmed that depolymerization occurs specifically at Lmod2-anchored PEs, not at distal, CP-capped BEs (Fig. 3E, movie S10). This behavior parallels profilin-induced depolymerization at formin-bound BEs^42,43^, suggesting that profilin can promote dissociation of monomers from both filament ends when elongation factors are bound. These results help explain the reduction in elongation rates observed at higher profilin concentrations (Fig. 3C).

In striated muscle, thin filaments are decorated with Tpm, which stabilizes filaments and regulates myosin binding^44^. Lmod2 is proposed to associate with Tpm at the PE through its TMBS^20^ (Fig. 1A), and Tpm enhances its nucleation activity^45^. To investigate whether Tpm also modulates Lmod2-dependent elongation, we measured the elongation rate of filaments anchored via Lmod2 with their BEs capped by CP across a range of concentrations of Tpm1.1 (Tpm’s predominant muscle isoform) and a fixed G-actin concentration. While Tpm1.1 reduced elongation rates in a concentration-dependent, biphasic manner (Fig. 3G), elongation persisted at the physiological Tpm1.1:actin ratio characteristic of muscle thin filaments (i.e., 1:7). The elongation rate plateau at higher Tpm concentrations likely reflects saturation of Tpm’s binding to filament sides and/or Lmod. While the underlying mechanism for this Tpm-dependent reduction in elongation is currently unclear, future experiments using Lmod2 variants lacking the Tpm-binding site may help resolve this question.

### Role of Lmod2 domains in processive PE elongation

Lmod2 contains three actin-binding sites (ABS1, LRR and WH2) (Fig. 1A)^21,45^, prompting us to investigate which of these domains might play a role in elongation. To address this, we measured elongation rates of filaments nucleated by coverslip-anchored wild-type (WT) as well as ABS1, WH2 and LRR mutants of human Lmod2 (Fig. 4A,C) in the presence of G-actin and CP.

**Fig. 4:**
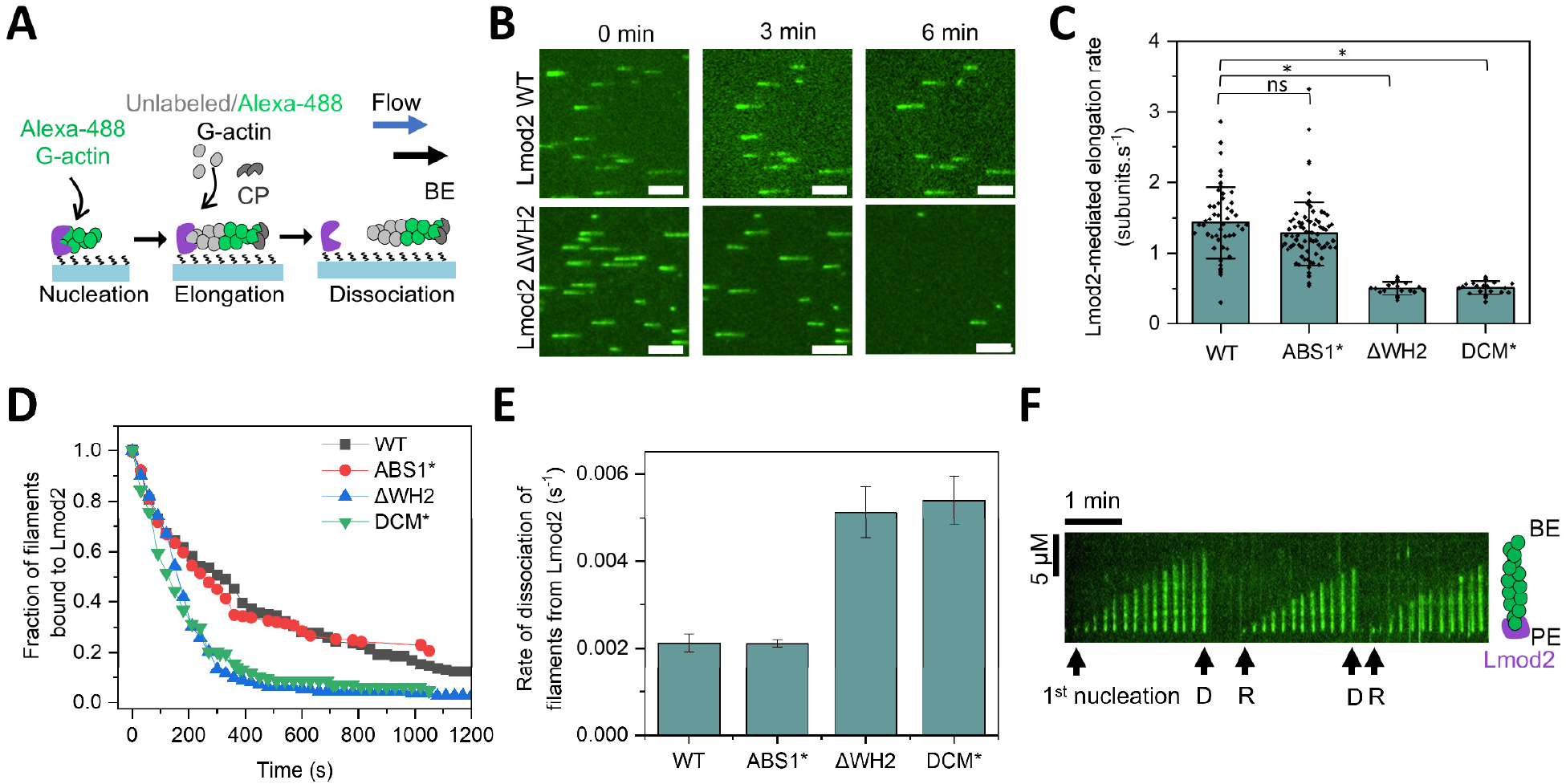
Role of Lmod2 domains in processive PE elongation. **(A)** Experimental strategy. Filaments were nucleated by coverslip-anchored Lmod2 using 1 µM Alexa-488–labeled G-actin (green), then elongated with 2 µM unlabeled G-actin plus 25 nM CP. PE elongation rates and the time-dependent survival fraction of Lmod2-bound filaments were measured. **(B)** Representative time-lapse images of filaments (green) detaching from surface-anchored Lmod2 WT (top) and the Lmod2ΔWH2 (bottom). Scale bar, 5 µm. **(C)** PE elongation rates of filaments nucleated by Lmod2 WT and mutants. Numbers of filaments analyzed: WT, 53; ΔWH2, 23; DCM*, 25; Linker*, 80. Statistical comparison by two-sample t-test against WT (ns, p ≥ 0.05; *, p < 0.05). **(D)** Survival fraction of Lmod2-bound filaments over time. Experimental data (symbols) were fit to single-exponential decays (lines) to obtain dissociation rate constants. Numbers of filaments: WT, 140; ΔWH2, 112; DCM*, 128; Linker, 175. **(E)** Dissociation rate constants of filaments nucleated by Lmod2 WT and mutants. Error bars indicate 65% confidence intervals from the exponential fits (see Methods). **(F)** Representative kymograph showing a filament re-nucleating at a previous detachment site from coverslip-anchored Lmod2 in the presence of 1 µM Alexa-488 G-actin (Movie S12). D, detachment and R, renucleation.

Point mutations in the region corresponding to Tmod1’s ABS1 (Fig. 1A) had no effect on Lmod2-mediated elongation rates (Fig. 4C), suggesting that this region does not participate in PE elongation *in vitro*. Similarly, it was also shown that ABS1 is dispensable for Lmod2-mediated nucleation^45^. In contrast, ABS1 is required for PE capping by Tmod, underscoring a functional divergence between the two proteins within this region. Consistent with previous results^21,45^, an LRR mutant (LRR*, Fig. 1A) failed to nucleate polymerization, precluding further analysis of elongation. In contrast, a WH2 deletion mutant (ΔWH2) retained the ability to nucleate and elongate filaments, but did so at less than 50% of the elongation rate observed with WT Lmod2 (0.64 ± 0.06 vs. 1.42 ± 0.50 subunits·s^−1^ at 2 µM G-actin; Fig. 4C).

We next assessed the processivity of Lmod2 using an approach previously applied to the formin mDia1^46-48^, in which the time-dependent detachment of Lmod2-nucleated filaments is monitored (Fig. 4A,B,D,E). Filaments nucleated by ΔWH2 dissociated ∼2-fold faster than those nucleated by WT Lmod2 (dwell time ∼195 s vs. ∼470 s; Fig. 4D,E, movie S11), corresponding to an ∼60% reduction in processivity. To exclude the possibility that this increased detachment rate arose from impaired coverslip attachment of ΔWH2 rather than faster PE dissociation, we performed re-nucleation experiments. Following filament dissociation, repeated filament nucleation and elongation were observed from the same spots. Comparable re-nucleation rates were observed at the initial Lmod2 locations for WT (44.5%) and ΔWH2 (45.4%) (Fig. 4F, fig. S5, movie S12), indicating that the faster detachment of ΔWH2-nucleated filaments reflects reduced PE processivity rather than compromised coverslip attachment.

Similar results were obtained with the DCM* mutant (Fig. 1A), which mimics the dilated cardiomyopathy–causing human *LMOD2* variant W398* (Fig. 4C,D,E). This mutation introduces a premature stop codon, generating a 45 kDa protein lacking both the PRD and WH2 domains (Fig. 1A). In humans, this Lmod2 variant causes early-onset of severe pediatric dilated cardiomyopathy^23,29^. Filaments formed by the DCM* and ΔWH2 mutants were ∼4-fold shorter than those formed by WT Lmod2, reflecting both their faster detachment and reduced elongation rate. Together, these findings provide a mechanistic explanation for the disease and are consistent with cellular studies showing that deletion of the WH2 domain leads to significantly shorter thin filaments in cardiomyocytes^19^.

## DISCUSSION

Over the past two decades, cellular and animal model studies have hinted at the existence of PE elongation in muscle thin filaments^13,16,49^. However, in the absence of a known PE polymerase, this mode of actin assembly has remained controversial. Here, we show Leiomodin-2 (Lmod2), a protein known to localize near the PEs of actin filaments in muscle sarcomeres^21^, acts as a processive elongator of actin filaments at their PEs. Using single-filament and single-molecule imaging, we show that Lmod2 remains persistently associated with elongating PEs and supports filament growth both from free and from profilin-bound actin monomers, indicating that Lmod2 can function under physiological conditions where the majority of ATP–actin is sequestered by profilin^3^.

Previous biochemical studies had suggested that Lmod functions exclusively as a nucleator in vitro^21^. However, Lmod’s nucleation function could not explain why Lmod depletion led to shorter thin filaments and its overexpression resulted in abnormally long thin filaments in cardiomyocytes^26,27^. Consequently, a discrepancy has persisted between Lmod’s known biochemical function and its effects *in vivo*^22-27^. Our results now provide a mechanistic explanation for these Lmod-related *in vivo* observations by demonstrating that Lmod2 directly promotes PE growth. Further analysis of Lmod2-mediated elongation indicates that PE growth likely occurs through a simple two-step kinetic process. At low actin concentrations, elongation rates scale with monomer concentration, whereas at higher concentrations elongation saturates, suggesting the presence of an Lmod2-dependent rate-limiting step. Although the exact structural basis of this step remains unresolved, our data are consistent with a model in which PE–bound Lmod2 alternates between monomer delivery and a subsequent rearrangement or stepping event. In this model, elongation is limited not by monomer availability but by Lmod2-dependent transitions at the filament end.

Compared to BEs, elongation at free PEs is inherently slow^50-52^. Surprisingly, we observe that Lmod2-bound PEs elongate faster than free PEs at low actin concentrations (<2 µM), including below PE critical concentration where free PEs depolymerize. Quantitative analysis indicates that this enhancement does not arise from an increased rate of monomer association, but instead from a strong suppression of monomer dissociation from the PE. Thus, unlike formins, which accelerate elongation primarily by increasing the effective on-rate of profilin–actin, Lmod2 enhances net growth by stabilizing terminal subunits at the PE. A recent cryo-EM study showed that the LRR domain of tropomodulin simultaneously interacts with the terminal three actin subunits at the PE^50^. Based on this, we propose that Lmod2 might achieve PE stabilization through a similar interaction of its LRR domain with terminal actin subunits. Such binding would prevent subunit dissociation and thereby enhance net PE elongation as observed here (Fig. 1G).

Lmod2-mediated polymerization shares important parallels with formin-mediated assembly. Both proteins can nucleate actin filaments and remain processively associated with filament ends following nucleation, with the key distinction that formins track BEs whereas Lmod2 tracks PEs. Despite these similarities, the underlying mechanisms differ fundamentally. Although both proteins can elongate from profilin-actin, formin-mediated elongation relies on its proline-rich domain (PRD) to recruit profilin–actin and increase the effective on-rate of monomer addition, whereas the PRD of Lmod2 is dispensable for elongation from profilin-bound monomers and Lmod-mediated elongation rate is slowed by profilin. Further, Lmod2-mediated elongation saturates at relatively lower monomer concentrations compared to formins, suggesting a mechanism that is limited by polymerase-dependent stepping rather than monomer recruitment. Together, these differences indicate that although Lmod2 and formins are both processive polymerases, they employ distinct strategies to promote actin filament elongation at opposite filament ends.

Lmod shares substantial domain architecture with tropomodulin (Tmod), including ABS1, the LRR domain, and tropomyosin-binding sites^53^. Despite these similarities, Lmod and Tmod have distinct effects at pointed ends: Lmod promotes pointed-end elongation, whereas Tmod enforces pointed-end capping. As described above, the LRR domain likely participates in stabilizing pointed end by Lmod (and preventing subunit dissociation) and Tmod (in preventing subunit association and dissociation). Despite this shared stabilizing function, Lmod and Tmod differ in their effects on elongation. In Tmod, ABS1 plays a critical role in pointed-end capping by binding the terminal actin subunit and sterically preventing further monomer addition^45,50^. In contrast, our results indicate that the homologous ABS1 domain in Lmod2 does not engage the pointed end in a capping configuration. Consistent with this, a previous study using isothermal titration calorimetry showed that ABS1 of Lmod2 does not interact detectably with actin^45^. As a result, Lmod2 stabilizes the pointed end against depolymerization while still permitting elongation. Together, these observations suggest that although Lmod and Tmod share a common mechanism for suppressing pointed-end disassembly, divergence in the conventional ABS1–actin engagement determines whether elongation is blocked, as in Tmod, or allowed, as in Lmod.

Beyond the LRR and ABS1 domains, Lmod2 also contains a C-terminal WH2 domain that binds actin monomers^53^. WH2 domains have classically been viewed as actin monomer–binding motifs that promote filament nucleation and/or severing by a number of actin-binding proteins, including Spire, Cordon Bleu, Leiomodin, VopF/VopL, and Sca2^54^. Our recent work has also suggested a role for WH2 domain in barbed-end depolymerization by CAP^55^ and pointed-end elongation by bacterial toxin VopF^56^. Although the WH2 domain is not strictly required for Lmod2-mediated PE elongation, its deletion markedly reduces elongation rates and filament dwell times, resulting in shorter filaments. Our biochemical findings are consistent with prior cellular studies in which deletion of the WH2 domain led to shortened thin filaments, supporting a role for WH2 in PE elongation *in vivo*^19^. While the reduction in dwell times in WH2-deletion mutants suggests a role for the WH2 domain in stabilizing Lmod2–filament interactions, it remains unclear whether WH2 domain also contributes to monomer delivery at the PE. Such a role would be conceptually analogous to the function of PRD domains in formins.

The existence of PE elongation in vertebrates raises important questions about how this activity is influenced by other PE regulators such as capping proteins like Tmod and depolymerases like CAP. Recent work from our lab and others has shown that BE dynamics are shaped by multicomponent interactions among polymerases, cappers, and depolymerases, including formin, capping protein, and twinfilin^46,55,57,58^. Whether analogous multicomponent regulation occurs at PEs, potentially involving Lmod, Tmod, and CAP, remains an important open question.

Finally, as recently proposed^13^, we expect that PE elongation will also have potential consequences for filament disassembly. Usually, actin filaments are thought to be rich in ATP-actin at their BEs and ADP-actin at their PEs. Lmod-mediated elongation would however generate filaments containing ATP–actin at both ends, with ADP–actin enriched in the filament interior. Such a nucleotide distribution could bias cofilin binding toward central regions of filaments while leaving filament ends relatively cofilin-free, potentially influencing severing and overall actin turnover dynamics.

### Conclusions and working model for Lmod2-mediated elongation of thin filaments

Our findings establish Leiomodin (Lmod) as the first vertebrate protein identified that is capable of processively elongating actin filaments at their PEs. Notably, Lmod2 promotes elongation in the presence of profilin, a condition that closely reflects the cytosolic environment, supporting the physiological relevance of this mechanism. By integrating our biochemical results with our previously published cellular and animal model studies, we propose a working model for how Lmod2 contributes to regulation of thin filament length in striated muscle (Fig. 5). During early stages of myofibrillogenesis, short nascent actin thin filaments, nucleated either spontaneously or by other nucleators like CapZ^59^, are capped at their BEs and anchored at the Z-disc. At this stage, tropomodulin-1 (Tmod1), which is expressed early in development^60^, binds PEs and enforces stable capping, effectively preventing both elongation and depolymerization^61^. Tmod1’s association with the PE is further stabilized through direct interactions with tropomyosin^62^.

**Fig. 5.**
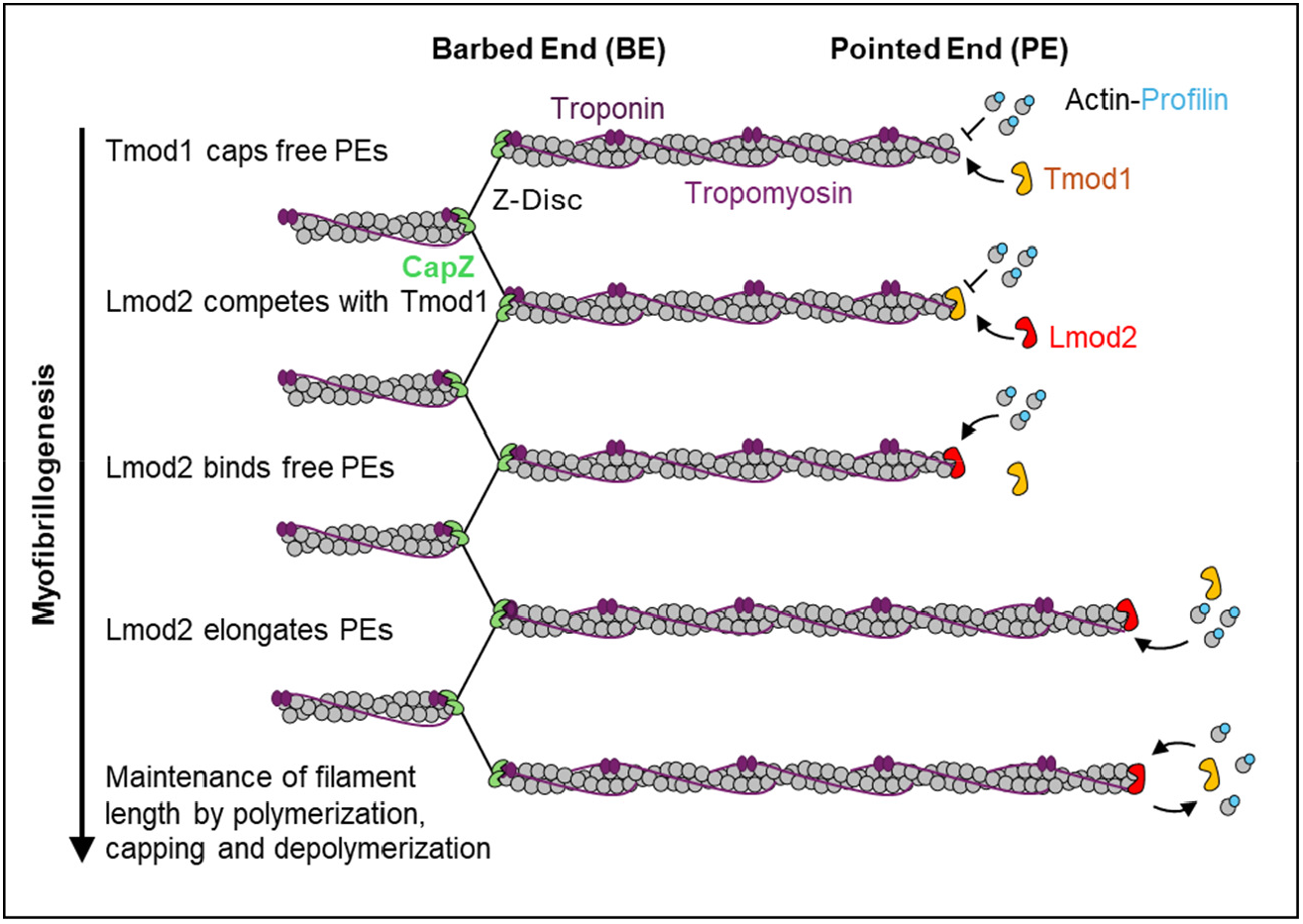
The working model for the role of Lmod2-mediated PE elongation in myofibrillogenesis. (i) A short nascent thin filament is nucleated by CapZ. Following nucleation, filament remains capped at its BE by CapZ and gets anchored in the Z disc via alpha-actinin. The filament PE gets capped by Tmod1. (ii) Lmod2 competes with Tmod1 for PE binding. (iii) Lmod2 processively elongates the actin filament from its PE by polymerizing profilin-bound G-actin while protecting it from Tmod1. (iv) In a mature filament, precise length is maintained by the dynamic control of polymerization (via Lmod2), capping (via Tmod) and depolymerization at the PE (depolymerization not shown).

As development progresses, Lmod2, which is expressed later than Tmod1^26^, competes with Tmod1 for PE binding. Upon binding, Lmod2 promotes polymerization from profilin-bound actin monomers. In mature thin filaments, we propose that filament length is maintained through a regulated balance between three processes: Lmod2-driven PE polymerization, Tmod1-mediated capping, and depolymerization mediated by actin turnover factors such as cyclase-associated protein and cofilin. This coordinated regulation would enable precise control of thin filament length, ensuring appropriate overlap with thick filaments for efficient sarcomeric contraction.

Dysregulation of thin filament length is associated with cardiac and skeletal muscle diseases, including dilated cardiomyopathy. In this context, our biochemical characterization of a well-established human LMOD2 truncation variant (c.1193G>A, p.Trp398*) is particularly informative^22,23,29,63^. Removal of the C-terminal WH2-containing region in this disease-associated variant results in a substantial reduction in elongation rate and processivity. These findings provide a mechanistic framework for understanding how impairment of Lmod2-mediated PE elongation may contribute to shortening of actin filaments observed in disease states *in vivo*.

Taken together, our work defines Lmod2 as a distinct class of vertebrate actin regulators that enables PE growth and reshapes actin filament dynamics. Our study provides a foundation for future investigations into how PE elongation, capping, and turnover are coordinated in cells.

## METHODS

### Purification and labeling of actin

Rabbit skeletal muscle actin was purified from acetone powder generated from frozen ground hind-leg muscle tissue of young rabbits (Pel-Freez). Lyophilized acetone powder stored at −80 °C was mechanically sheared in a coffee grinder, resuspended in G-buffer (5 mM Tris-HCl, pH 7.8, 0.5 mM dithiothreitol, 0.2 mM ATP, and 0.1 mM CaCl_2_), and cleared by centrifugation for 20 min at 50,000 × *g*. The supernatant was collected and filtered through Whatman paper. Actin was polymerized overnight at 4 °C with slow stirring by addition of 2 mM MgCl_2_ and 50 mM NaCl. NaCl was then added to a final concentration of 0.6 M, and stirring was continued for an additional 30 min at 4 °C. Actin filaments were pelleted by centrifugation for 150 min at 280,000 × *g*, resuspended by Dounce homogenization, and dialyzed against G-buffer for 48 h at 4 °C. Monomeric actin was precleared at 435,000 × *g* and loaded onto a Sephacryl S-200 16/60 gel-filtration column equilibrated in G-buffer. Actin-containing fractions were stored at 4 °C and used within four weeks of purification.

For fluorescent labeling, G-actin was polymerized by overnight dialysis against modified F-buffer (20 mM PIPES, pH 6.9, 0.2 mM CaCl_2_, 0.2 mM ATP, and 100 mM KCl). Actin filaments were incubated for 2 h at room temperature with a fivefold molar excess of Alexa Fluor 488 NHS ester (Thermo Fisher Scientific). Labeled actin filaments were pelleted by centrifugation at 450,000 × *g* for 40 min, resuspended in G-buffer, homogenized with a Dounce, and incubated on ice for 2 h to depolymerize filaments. Actin was re-polymerized by addition of 100 mM KCl and 1 mM MgCl_2_, pelleted again by centrifugation, homogenized, and dialyzed overnight at 4 °C against G-buffer. The solution was clarified by centrifugation, and actin concentration and labeling efficiency were determined.

### Purification of profilin

Human profilin-1 was expressed in *E. coli* strain BL21 (pRare) grown to log phase in LB broth at 37 °C and induced with 1 mM IPTG for 3 h at 37 °C. Cells were harvested by centrifugation at 15,000 × *g* at 4 °C and stored at −80 °C. For purification, cell pellets were thawed and resuspended in 30 mL lysis buffer (50 mM Tris-HCl, pH 8.0, 1 mM DTT, 1 mM PMSF, and protease inhibitors consisting of 0.5 µM each of pepstatin A, antipain, leupeptin, aprotinin, and chymostatin). Cells were lysed by tip sonication on ice. The lysate was clarified by centrifugation for 45 min at 120,000 × *g* at 4 °C. The supernatant was applied to 20 mL poly-L-proline–conjugated beads packed in a disposable column. Beads were washed at room temperature with wash buffer (10 mM Tris, pH 8.0, 150 mM NaCl, 1 mM EDTA, and 1 mM DTT), followed by two column volumes of the same buffer containing 3 M urea. Profilin was eluted with five column volumes of buffer containing 8 M urea. Pooled fractions were concentrated and dialyzed against 4 L of dialysis buffer (2 mM Tris, pH 8.0, 0.2 mM EGTA, 1 mM DTT, and 0.01% NaN_3_) for 4 h at 4 °C. The dialysis buffer was replaced and dialysis was continued overnight at 4 °C. Following dialysis, the protein was clarified by centrifugation for 45 min at 450,000 × *g* at 4 °C.

### Purification, biotinylation and labeling of capping protein

Mouse His-tagged capping protein (CP) was expressed in *E. coli* BL21(DE3) pLysS cells. CP subunits α1 and β2 were expressed from the same plasmid, with a single His tag on the α subunit, as done previously^47^. SNAP-tagged CP (SNAP-CP) was expressed from a single plasmid encoding a His- and SNAP-tagged β1 subunit and an untagged α1 subunit, as done previously^58^. Purified SNAP-CP was incubated overnight at 4 °C with a fivefold molar excess of SNAP-Surface 549 dye (New England Biolabs). Free dye was removed using PD-10 desalting columns (Cytiva). For biotin labeling, purified SNAP-CP was incubated overnight at 4 °C with a twofold molar excess of SNAP-biotin (New England Biolabs). Free biotin was removed using PD-10 desalting columns.

### Purification and labeling of Lmod2

Human and mouse Leiomodin 2 (Lmod2; wild-type and mutant proteins) were expressed and purified as described previously^45^. For biotin labeling, purified SNAP-Lmod2 was incubated overnight at 4 °C with a twofold molar excess of SNAP-biotin (New England Biolabs). Free biotin was removed using PD-10 desalting columns (Cytiva). For dye labeling, purified SNAP-Lmod2 was incubated overnight at 4 °C with a fivefold molar excess of SNAP-Surface 549 dye (New England Biolabs). Free dye was removed using PD-10 desalting columns.

### Microfluidics-assisted TIRF (mf-TIRF) imaging and analysis

For all mf-TIRF experiments^36^, coverslips were first cleaned by sonication in Micro90 detergent for 20 min, followed by successive 20 min sonications in 1 M KOH, 1 M HCl and 200 proof ethanol for 20 min each. Washed coverslips were stored in fresh 200 proof ethanol. Prior to each experiment, coverslips were washed extensively with H_2_O and dried in an N_2_ stream. These dried coverslips were coated with 2 mg/mL methoxy-poly(ethylene glycol) (mPEG)-silane (MW 2,000) and 2 µg/mL biotin-PEG-silane (MW 3,400; Laysan Bio, USA) in 80% ethanol (pH 2.0) and incubated overnight at 70 °C. A 40 µm-high PDMS mold with three inlets and one outlet was mechanically clamped onto a PEG-silane–coated coverslip. The chamber was connected to a Maesflo microfluidic flow-control system (Fluigent, France), rinsed with mf-TIRF buffer (10 mM imidazole, pH 7.4, 50 mM KCl, 1 mM MgCl_2_, 1 mM EGTA, 0.2 mM ATP, 10 mM DTT, 1 mM DABCO), and incubated with 1% BSA and 10 µg/mL streptavidin in 20 mM HEPES (pH 7.5) and 50 mM KCl for 5 min. Coverslips were subsequently functionalized with Lmod2 or CP as needed for specific experiments.

### Image acquisition and analysis

Single-wavelength time-lapse TIRF imaging was performed on a Nikon Ti2000 inverted microscope equipped with 488-nm and 561-nm laser lines, a 60× TIRF objective (numerical aperture 1.49), and an EMCCD camera (iXon Life 888, Andor). The pixel size was 144 × 144 nm. Focus was maintained using a hardware autofocus system. Time-lapse images were acquired using Nikon Elements software. For two-color imaging, samples were sequentially excited with 488-nm and 561-nm lasers.

Images were analyzed using Fiji. Background subtraction was performed using a rolling-ball algorithm (radius, 5 pixels), and drift correction was applied using the Image Stabilizer plugin. For each condition, filaments were acquired from multiple fields of view.

For elongation rate measurements, kymographs of individual filaments were generated using the built-in kymograph tool, and elongation rates were calculated from kymograph slopes assuming a monomer length of 2.7 nm per actin subunit.

For filament detachment measurements (Fig. 2I), time-lapse images were used to determine survival fraction data, which were fit with a single-exponential decay to obtain dissociation rate constants for wild-type and mutant Lmod2. Detachment events were identified from kymographs.

Data analysis and curve fitting were performed using Microcal Origin. All experiments were repeated at least three times with similar results. Representative datasets are shown. Data are reported as mean ± s.d. Statistical comparisons between two conditions were performed using two-sided Student’s *t* tests. Kinetic parameters were obtained by fitting the data to linear or a saturating binding curve as indicated, and statistical significance was defined as *P* < 0.05.

## Acknowledgements

We thank Drs. Alla Kostyukova and Christopher Pappas for assistance with designing the Lmod2 mutants, Dr. Mert Colpan for initiating the collaboration, and Surbhi Garg for preliminary experiments. We also thank Dr. Dimitrios Vavylonis for helpful discussions. We thank Shoichiro Ono for comments on the manuscript. This work was supported by NIH R35GM143050 (S.S.), R01GM120137 (C.C.G) and R01HL123078 (C.C.G. and S.S.).

## Author contributions

S.S. and C.C.G. conceptualized the study. S.B. conducted experiments. T.L. generated key reagents. S.C. helped with data analysis. S.S. supervised the research and wrote the manuscript. All authors contributed to methodology, investigation, visualization and editing of the manuscript.

## Competing interests

We declare no conflicts of interest.

## Notes

### Competing Interest Statement

The authors have declared no competing interest.

